# Multi-objective formulation of MSA for phylogeny estimation

**DOI:** 10.1101/418095

**Authors:** Muhammad Ali Nayeem, Md. Shamsuzzoha Bayzid, Atif Hasan Rahman, Rifat Shahriyar, M. Sohel Rahman

**Author notes:** Corresponding author. Professor, Dept. of CSE, BUET, Dhaka 1000, Bangladesh.

## Abstract

Multiple sequence alignment (MSA) is a basic step in many analyses in computational biology, including predicting the structure and function of proteins, orthology prediction and estimating phylogenies. The objective of MSA is to infer the homology among the sequences of chosen species. Commonly, the MSAs are inferred by optimizing a single function or objective. The alignments estimated under one criterion may be different to the alignments generated by other criteria, inferring discordant homologies and thus leading to different evolutionary histories relating the sequences. In recent past, researchers have advocated for the multi-objective formulation of MSA, to address this issue, where multiple conflicting objective functions are being optimized simultaneously to generate a set of alignments. However, no theoretical or empirical justification with respect to a real-life application has been shown for a particular multi-objective formulation. In this study, we investigate the impact of multi-objective formulation in the context of phylogenetic tree estimation. Employing multi-objective metaheuristics, we demonstrate that trees estimated on the alignments generated by multi-objective formulation are substantially better than the trees estimated by the state-of-the-art MSA tools, including PASTA, MUSCLE, CLUSTAL, MAFFT etc. We also demonstrate that highly accurate alignments with respect to popular measures like sum-of-pair (SP) score and total-column (TC) score do not necessarily lead to highly accurate phylogenetic trees. Thus in essence we ask the question whether a phylogeny-aware metric can guide us in choosing appropriate multi-objective formulations that can result in better phylogeny estimation. And we answer the question affirmatively through carefully designed extensive empirical study. As a by-product we also suggest a methodology for primary selection of a set of objective functions for a multi-objective formulation based on the association with the resulting phylogenetic tree.

## 1 Introduction

In biological research, multiple sequence alignment (MSA) is a useful and/or essential task in various applications such as phylogeny estimation, prediction of the structure and function of a RNA or protein, identification of functionally important sites, orthologous gene identification etc. The MSA task seeks to arrange more than two biological sequences based on certain criteria (such as evolutionary history, 3D structure etc.) by inserting spaces between letters in the sequences. In this research, we limit our focus on MSA in the context of phylogeny. Phylogeny estimation from molecular sequences generally operates as a two-phase approach. At first the given sequences are aligned using a MSA method, and then a tree is estimated from the resultant alignment. The quality of inferred trees heavily depends on the quality of the corresponding alignment. Therefore, it is important to select a MSA tool that is the ‘most suitable’ in the phylogenetic context.

In this study, we make an attempt to design a multi-objective formulation of MSA that is more effective in phylogeny estimation. Our motivation for a multi-objective formulation comes from the fact that the alignment estimated under one objective may be different to the alignments generated by other objectives, inferring discordant homologies and thus leading to different and often conflicting evolutionary histories relating the sequences under consideration. Multi-objective formulations can address this issue by optimizing multiple conflicting objectives simultaneously to generate a set of alignments. However, we are faced with the challenge of using appropriate measures/metrics to choose from among a number of objective sets to optimize. So, we ask the natural question whether the popular general purpose measures to judge the alignment quality can truly reflect the quality in the context of a particular application domain, i.e., phylogeny estimation in our case. While this question has received some shallow discussion in several studies [1, 2, 3], to the best of our knowledge no systematic investigation has been reported in the literature to this end. Therefore, in essence, we systematically investigate whether a phylogeny-aware metric can guide us better in choosing appropriate multi-objective formulation or tools capable of generating alignments that can produce better phylogenetic trees.

There are numerous tools available in the literature to compute MSA. We can broadly divide them into three groups: progressive techniques, consistency based techniques and iterative techniques. This division is not exclusive as many tools also use combination of these techniques. Progressive technique is the foundation of many MSA tools such as, Clustal Ω [4], PRANK [5], Kalign [6], FSA [7], RetAlign [8] etc. They compute the alignment using a guide tree by aligning pairs of sequences in a “bottom-up” manner. The consistency based techniques first construct a database of local and global pairwise alignments to facilitate generating an overall accurate alignment. The representatives of this category are T-Coffee [9], ProbCons [10], MSAProb [11], ProbAlign [12] etc. On the other hand, the iterative techniques were designed to achieve reliable alignments. These techniques try to fix the effect of mistakes made during the initial phases by repeating some crucial steps until some criteria are met. We find several examples of such techniques, such as, MAFFT [13], MUSCLE [14], MUMMALS [15], ProbCons, PRIME [16], SAGA [17] etc. In this category, we also see some “meta-methods” such as, SATé [3] and PASTA [2], which co-estimate alignment and tree using other methods. These tools achieve scalability by employing the divide-and-conquer principle and are being used widely in practice.

The performance of an MSA tool is usually evaluated by comparing its output alignment with the reference alignment (provided with the dataset as the ground truth) in terms of several measures. To this end, the most popular measures are perhaps sum-of-pair (SP) score and total-column (TC) score. SP score is the fraction of the homologies (i.e., pairs of aligned characters) in the reference alignments recovered in the estimated alignment. Similarly TC score is the fraction of the actual aligned columns that appears in the estimated alignment.

In this post-genomic era, the MSA datasets are posing new challenges to the researchers. Usually an MSA method is provided with a default parameter configuration for aligning any problem instance with a satisfactory accuracy. But we know that these default values can not guarantee the best output throughout all kinds of datasets [18]. For instance, there is a parameter in ProbCons called the number of iterative refinement passes. Although it can vary between 0 to 1000 the default value is set to 100. We can achieve better results by tuning the parameter values which is not a straightforward task. Moreover, despite rigorous parameter tuning, no method can consistently outperform other methods for all datasets.

Therefore, we see the emergence of novel approaches that combine different alignment tools [19]. One such approach is metaheuristics where alignments generated from different tools are exploited to produce improved alignments without vesting any effort in parameter tuning. The success of such a metaheuristic approach depends on the selection of proper objective function that can push the solutions (i.e., alignments) towards the desired zone that reflects the actual purpose of an alignment task. As any single objective function alone can not be effective to tackle different challenges, it is wise to simultaneously optimize multiple objective functions. This will produce a set of competing solutions as the final output, which can be expected to contain our desired solution(s). Thus the formulation of MSA as a multi-objective optimization problem turns out to be appealing.

During the last decade, we find several studies [20, 21, 22, 23, 24, 25, 26, 27] with multiobjective formulation for MSA have been published – proposing two to four objective functions to capture and quantify different aspects of an alignment. Among them probably the most popular is the sum-of-pairs score and its weighted variants, where pairwise score is calculated for each pair of aligned sequences using a substitution matrix. This matrix should reflect the characteristics of the data at hand. Although we know that the same character across all rows of a column does not necessarily indicate homology, the count of such columns in an alignment is seen as a maximization objective known as totally conserved columns. Next, we find attempts to minimize total number of gaps to maintain the compactness of an alignment. Then there are different types of gap penalties that penalize each sequence for introducing gaps. Also we find two other objective functions, Entropy and Similarity, that compute column-wise scores and then sum those together. Both of them try to express the homogeneity of characters in a column using two different ways. Contrary to the performance/quality measures mentioned earlier (such as SP score, TC score), we are not allowed to use the reference alignment while calculating these objective functions.

We notice several issues in the works advocating multi-objective formulation of MSA (in the context of different applications where the MSA will be used). First of all, in these works, there is a lack of sound theoretical or empirical justification for the choice of a particular objective function to be optimized. Secondly, we also notice the absence of a sound rationale/justification behind the two most popular performance metrics, namely, sum-of-pair score and total column score. On the contrary, it seems only natural that, performance score should reflect the actual purpose of MSA. For example, if the goal is to estimate a phylogenetic tree, the performance metric to be used for evaluation should be able to accurately measure the quality and usefulness of the constructed tree. Notably, another issue, specific to the domain of phylogeny estimation, is the use of relatively smaller (number of taxa below 50) datasets in experiments.

In this article, we attempt to demonstrate the effectiveness of multi-objective MSA by addressing the above mentioned limitations in the context of its intended application domain (i.e., phylogeny estimation). To make fair comparison with nine state-of-the-art MSA tools, we conduct comprehensive experimentations on both simulated and biological datasets using tree as well as alignment quality measures. Our study represents the only known work on devising a phylogeny-aware multi-objective formulation for MSA. In particular, this article makes the following key contributions:

- To the best of our knowledge, this is the first attempt to investigate the effect of using domain specific measures (as opposed to generic alignment measures) to evaluate the performance of MSA methods in the context of phylogeny estimation.
- We suggested a methodology based on multiple linear regression to judge the potential efficacy of a multi-objective formulation of MSA. Then, based on this methodology, we proposed two multi-objective formulations that had the potential to yield better phylogenetic trees.
- Finally, we demonstrated that the multiobjective formulations can consistently yield better phylogenetic trees than several state-of-the-art MSA tools. And interestingly we found that, popular alignment quality measures do not necessarily lead to highly accurate phylogenetic trees.

## 2 Results

We conducted extensive experiments with a 100- taxon simulated dataset, two biological rRNA datasets and 27 BAliBASE datasets. We begin by carefully and systematically selecting two multi-objective formulations which are potentially useful for phylogenetic tree estimation. Then for each dataset, we generate alignments through running multi-objective metaheuristics as well as nine state-of-the-art MSA tools. Then we compare those alignments with respect to both generic and domain specific quality measures. In this section, we discuss our obtained results after a brief discussion on our experimental design and datasets. In what follows, unless otherwise specified, when we discuss the (best) results of a tool, we mean one of the abovementioned nine tools.

### 2.1 Experimental design

Our experimental methodology is briefly described below (please see also Figure 1):

- <u>Step 1:</u> Following a systematic approach involving multiple linear regression applied on a simulated dataset, we first make an attempt to identify and choose two multiobjective formulations that turn out to be potentially more effective in the context of phylogeny estimation.
- <u>Step 2:</u> We run a popular and effective multi-objective metaheuristics on both biological and simulated datasets to optimize each set of objective functions selected in Step 1. Each run of the metaheuristics on each dataset gives us a set of alignments as output.
- <u>Step 3:</u> We also run nine state-of-the-art MSA tools (please see Section 4, Table 4) to generate alignments on all these datasets.
- <u>Step 4:</u> We evaluate the quality of each generated alignment with respect to the reference alignment using two popular measures, namely, SP score and TC score.
- <u>Step 5:</u> For each of the generated alignments, we infer maximum likelihood (ML) phylogenetic tree. Then we measure the quality of each inferred tree with respect to the reference tree (true tree) using the mostly used measure in the literature called false negative (FN) rate.
- <u>Step 6:</u> Finally we compare the alignments and the corresponding ML trees generated by the multi-objective optimization with the ones generated by the state-of-the-art tools.

**Figure 1:**
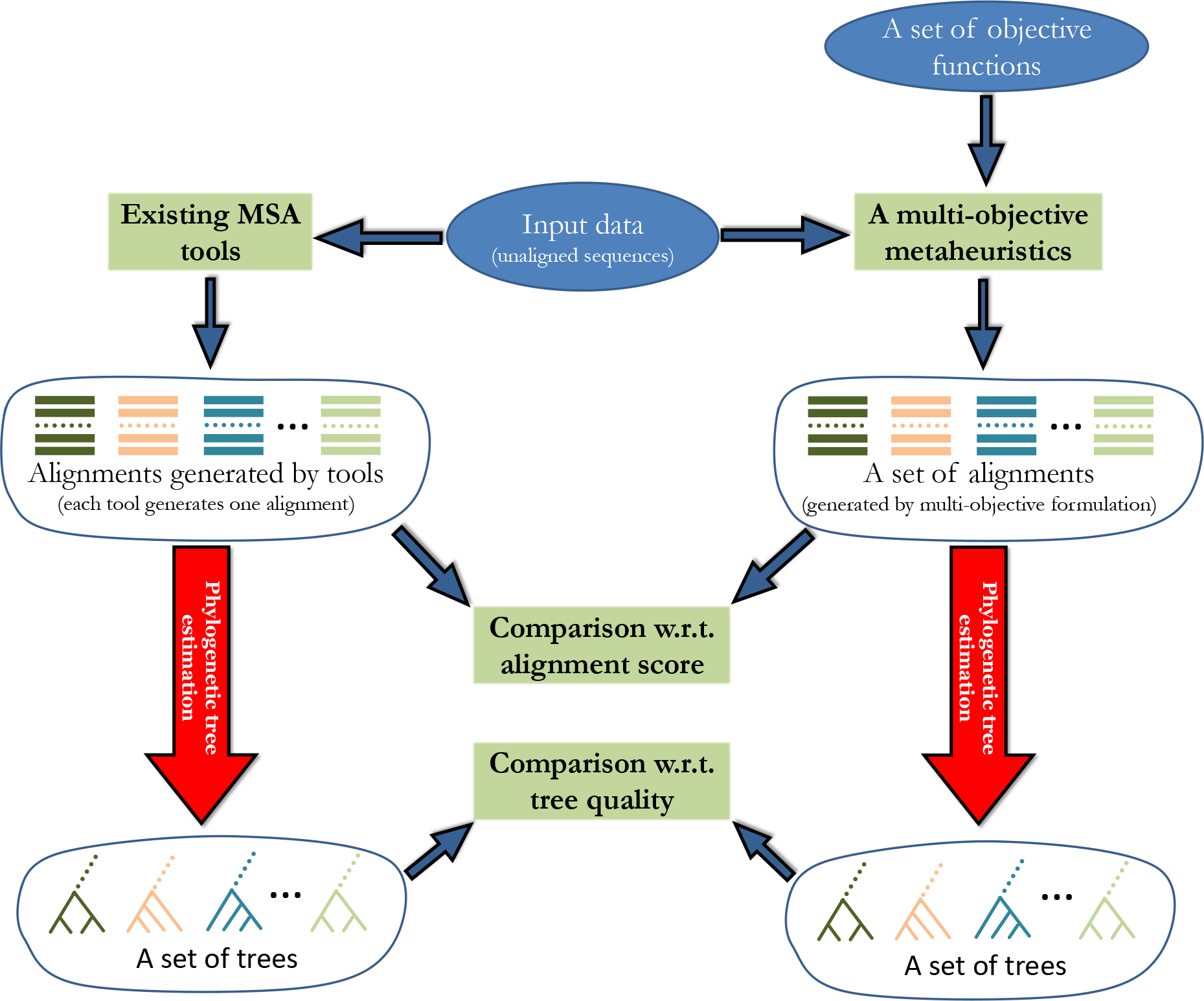
Our methodology for finding the impact of a multi-objective formulation (i.e., a set of objective functions) of MSA on phylogenetic tree estimation. For each dataset (i.e., unaligned sequences), we run an multi-objective metaheuristics. It simultaneously optimizes the given objective functions and outputs a set of alignments which represents the best-possible compromise among all objective functions. We also run several existing MSA tools on that dataset and each tool generates one alignment. We evaluate the quality of each generated alignments with respect to the reference alignments using widely used scores. Also we estimate phylogenetic trees for all alignments and evaluate each tree with respect to the reference. Then we compare the alignments and the corresponding phylogenetic trees generated by the multi-objective formulation with the ones generated by the existing tools based on alignment score as well as tree quality. We also observe the association between alignment scores and tree quality values to examine whether it is appropriate to use alignment score in the context of phylogeny estimation.

### 2.2 Datasets

We studied three datasets: 100-taxon simulated dataset, Biological rRNA datasets and BAliBASE datasets. We use the 100-taxon simulated dataset for dual purposes. At first, we conduct experiments with this dataset to examine whether a multi-objective formulation of MSA is potentially phylogeny-aware and in the sequel, we select two such formulations. Afterwards, we validate the effectiveness of these formulations against the state-of-the-art MSA tools based on this dataset as well as other biological datasets. We now introduce each of these datasets.

#### 2.2.1 100-taxon simulated dataset

We used 10 (out of 20) randomly selected replicates (R0, R2, R4, R6, R9, R10, R13, R14, R17, R19) of simulated nucleotide dataset from the study of 3. It is publicly available at https://sites.google.com/eng.ucsd.edu/datasets/sate-i. Table S1 in the supplementary file gives the reference alignment statistics for this dataset.

#### 2.2.2 Biological rRNA datasets

We analyzed two biological ribosomal RNA datasets, 23S.E and 23S.E.aa ag, from 3 which are challenging for phylogeny estimation methods. Each of these datasets is given with a highly reliable, curated reference alignment from Gutell Lab. The statistics of the reference alignments of these datasets are presented in Table S2 of the supplementary file. Reference trees for these datasets were generated from the reference alignments by running RAxML [28] with bootstrapping, and retaining only the highly supported edges. We evaluated generated alignments with respect to the reference alignment using the tool FastSP [29].

#### 2.2.3 BAliBASE datasets

BAliBASE 3.0 [30] is the most widely used benchmark alignment databases of protein families. It provides manually refined reference alignments of high quality based on 3D structural superposition. These datasets are organized into six groups according to their families and similarities: RV11 (very divergent sequences, residue identity below 20%), RV12 (medium to divergent sequences, 20%-40% residue identity), RV20 (families with one or more highly divergent sequences), RV30 (divergent subfamilies), RV40 (sequences with large terminal N/C extensions), and RV50 (sequences with large internal insertions). In this study, we selected four to five representative datasets from each group as reported in Table S3 of the supplementary file. We generated reference trees for these datasets by running RAxML with bootstrapping. We evaluated estimated alignments with respect to the core blocks (regions for which reliable alignments are known to exist) using the program bali score available at http://www.lbgi.fr/balibase/BalibaseDownload/.

### 2.3 Selection of multi-objective formulations

As has been mentioned above, we have used 100-taxon simulated dataset for dual purposes: to select one or more multi-objective formulations that have the potential to be ‘phylogeny-aware’ (Section 2.3.1, 2.3.2) and to judge the efficacy of the detected formulations in comparison with the other state-of-the-art tools (Section 2.4). We first conduct extensive experiments to choose a formulation (i.e., a set of objective functions) of MSA from among the existing popular ones from the literature (Section 2.3.1); subsequently, we also suggest a new promising formulation (Section 2.3.2).

#### 2.3.1 Selection from among the existing formulations

To reduce the computational effort, we preselect three multi-objective formulations of MSA and limit our investigation thereon. Thus we choose one of the formulations from among {Gap, SOP} [23], {SOP, TC} [20] and {wSOP, TC} [31, 26] (please see Section 4 Table 2). We experiment with five randomly selected replicates (R0, R4, R9, R14, R19) and then judge based on two criteria: firstly, we used multiple linear regression analysis to examine the association between individual objective function and FN rate; secondly, we assess the alignments generated through the optimization of each set of objective functions in terms resultant ML trees.

**Table 2:** Terms used to denote objective functions.

| # | Objective function | Term |
| --- | --- | --- |
| 1 | Maximize no. of totally aligned columns | TC |
| 2 | Minimize no. of gaps | Gap |
| 3 | Maximize sum of pairs | SOP |
| 4 | Maximize weighted sum of pairs with affine gap penalties | wSOP |
| 5 | Minimize entropy | Entropy |
| 6 | Maximize similarity based on gap containing columns | SimG |
| 7 | Maximize similarity based on non-gap columns | SimNG |
| 8 | Maximize concentration of gaps | GapCon |

We need to consider the relationship between each pair of objective functions to properly interpret the result of multiple linear regression. We perform this by running an appropriate multiobjective metaheuristic (i.e., NSGA-III [32]) for 25 times which optimizes all the objective functions (i.e., {Gap, SOP, wSOP, TC}) and thus we obtain a large collection of diverse alignments. A visualization of the interrelations among the objective values of those solutions is presented in Figure S2 of the supplementary file. From these experiments, we have the following two key observations.

a. In all the cases, SOP is totally correlated with wSOP. So we do not need to optimize both of them. Moreover, this high correlation creates a serious problem in multiple regression analysis called multicollinearity [33]. Therefore, we should not keep these two objective functions together in our regression analysis. Also it is redundant to considering both of them in the multiobjective formulation.
b. SOP is clearly in conflict with Gap across all the replicates. Therefore, if we optimize them simultaneously, we can generate many diverse solutions which represent the compromise between these two objective functions [34]. These diverse collection is likely to contain the desired alignment for any kind of dataset.

As the objective functions are inter-related, we need to measure the degree of association between an objective and FN rate while holding the remaining objectives constant to avoid getting any spurious result [33]. Therefore, we perform multiple linear regression by employing the following model:

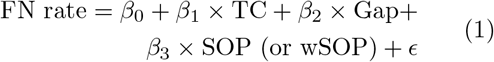

Each coefficient (*β*_1_, *β*_2_ and *β*_3_) represents the expected change in the FN rate per unit change in the corresponding objective function when all the remaining objective functions are held constant. For this reason they (*β*_*i*_) are called partial regression coefficients. *ϵ* is the random error component which is assumed to follow a Gaussian distribution with mean zero and some fixed standard deviation. We fit this model to the solutions generated by optimizing the set {Gap, SOP, wSOP, TC}. For each of those solutions, we estimate ML tree and evaluate its quality in terms of FN rate. We estimate these coefficients using least-squares method (an illustration is presented in Figure S3 of the supplementary file). We apply *t*-test on individual regression coefficient (i.e., slope) *β*_*i*_ (with null hypothesis *β*_*i*_ = 0) to test the significance of that association. We can note the following two interesting points from these results.

a. In majority of the cases (R0, R4 and R14), Gap, SOP and wSOP exhibit a good degree of association with FN rate (i.e positive slope) with high confidence (p-value close to 0) compared to other objective functions. So, we can expect them to be good optimization objectives for MSA.
b. For replicate R4 and R19, none of the objective exhibit good association. This shows that an objective function might not perform well across all problem instances.

Now we measure the strength of each objective set based on the FN rate achieved by the members of generated solution set. To accomplish this, For each set of objective functions, we run a suitable multi-objective metaheuristics (NSGAII [35]) for 20 times following the standard practice of operations research (OR) literature (due to the stochastic nature of metaheuristics). Each run generates a set of solutions that represents the trade-offs in satisfying all objectives. Afterwards, we inferred ML tree for each of the generated alignment. We collected the best FN rates from each of the 20 solution sets and examine the distribution of these FN rates (a visualization of these distributions using boxplots is presented in Figure S4 of the supplementary file). Here we have the following key observations:

- For most of the cases, the combined set {TC, Gap, SOP, wSOP} achieves better results than the other sets. This indicates that adding suitable objective functions increase the chance of achieving the best FN rate. However, this increases the overall complexity of the multi-objective metaheuristic. So in this study we keep the size of objective set as small as possible.
- Among our three pre-selected objective sets, {Gap, SOP} achieves relatively lower FN rates. This is consistent with our regression results discussed earlier.
- Both {TC, Gap, SOP, wSOP} and {Gap, SOP} persistently generate better FN rates than the state-of-the-art tools.

Based on our findings discussed so far, we consider {Gap, SOP} to be the most suitable candidate to conduct our study among all the formulations considered above.

#### 2.3.2 Selection of a new formulation

The results reported in the last subsection suggest that the objective functions that exhibit good association with FN rate should be more effective than the other objective functions for estimating phylogenetic trees. Based on this we make an attempt to form a new objective set as follows. We first propose four new objective functions that quantify different aspects of MSA: Entropy, SimG, SimNG and GapCon (the details are presented in Section 4.1). We combine these with TC and Gap and run NSGA-III to optimize the objective set {Entropy, TC, Gap, SimG, SimNG, GapCon} for 40 times to generate numerous diverse alignments. We used those to examine the association of our proposed objective functions with FN rate using multiple linear regression analysis (please refer to Figure S5 of the supplementary file for a visualization of the relationship between the relevant pairs of objective functions within the set). The key observations of this analysis are summarized as follows:

- Entropy has a strong correlation with SimG which is problematic for multiple regression analysis. So, we should not keep these two objectives at the same time in our regression model as well as in the multi-objective formulation.
- SimG and SimNG are in conflict with each other. So by optimizing them simultaneously, a multi-objective metaheuristic can generate large number of diverse alignments.

Now we express the relationship between FN rate and the proposed objective functions using the following model:

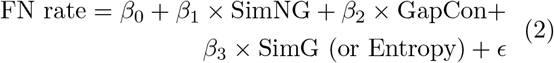

We estimate the regression coefficients by fitting the above model to the solutions generated by optimizing the objective set {Entropy, TC, Gap, SimG, SimNG, GapCon}. Here we find that, in each case, both SimG and SimNG exhibit positive correlation with FN rate. So we choose {SimG, SimNG} as our new objective set. For a visualization of the results using partial regression plots please refer to Figure S6 of the supplementary file.

### 2.4 Results on 100-taxon simulated dataset

We now examine the performance of the two objective sets selected (in the previous sections), namely, {Gap, SOP} and {SimG, SimNG} on randomly chosen 10 replicates of 100-taxon simulated dataset against the state-of-the-art tools. Following the standard practice in the OR literature, we conduct 20 runs of NSGA-II for each possible combination (replicate, set of objective functions) due to its (NSGA-II) stochastic nature. Each run produces a set of 100 alignments that represent the trade-offs in satisfying all objectives under consideration. We first evaluate each alignment using the widely used metrics, namely, TC and SP score. Next, we estimate ML tree for each alignment and measure the FN rate of the resultant tree. We use the averaged TC score, SP score and FN rate over 10 replicates to evaluate the strength of each objective set from two perspectives in Figure 2.

**Figure 2:**
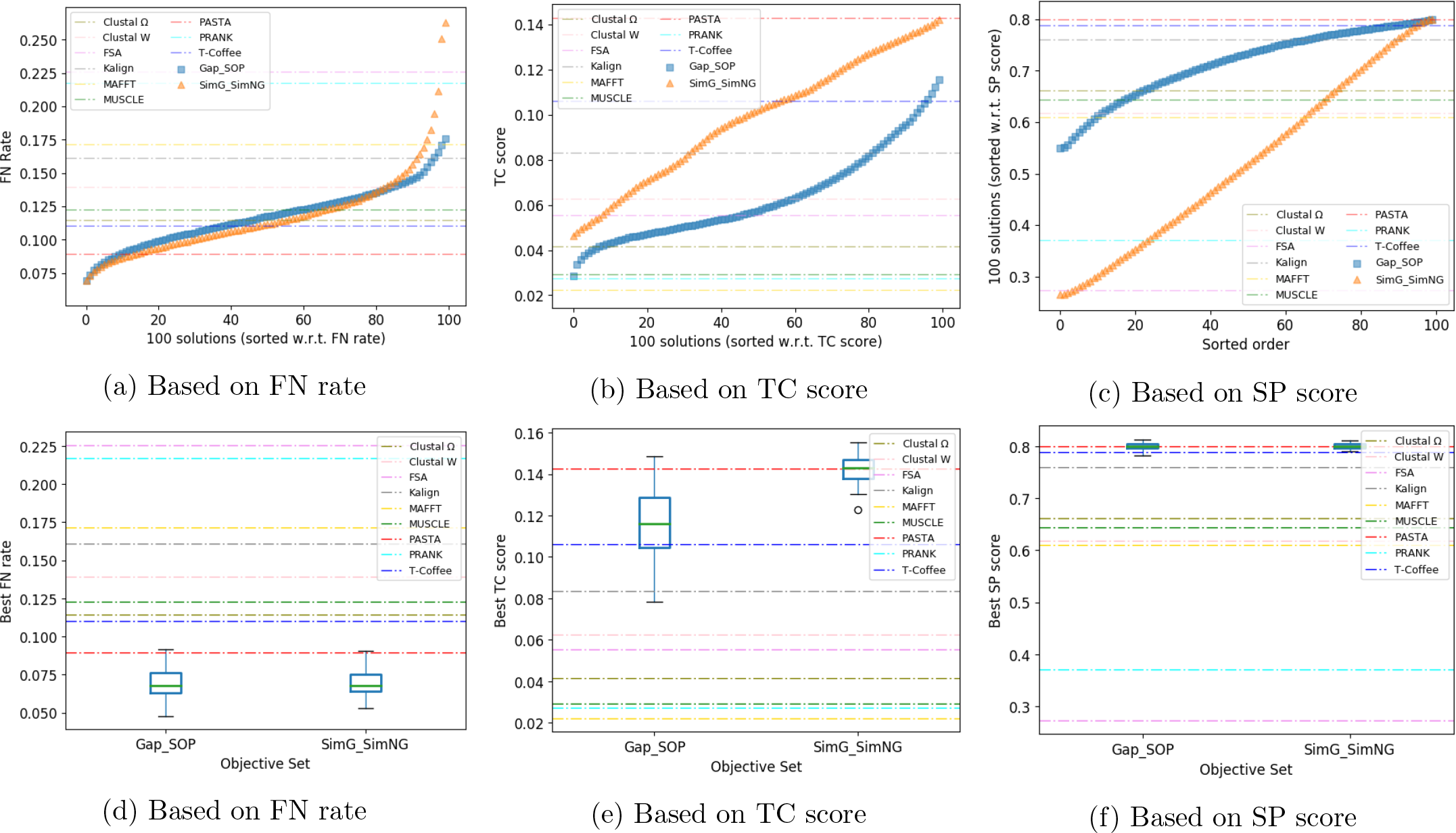
100-taxon simulated dataset: **Top panel** (part (a) - (c)) illustrates how the 100 solutions provided by the multi-objective formulations perform with respect to each of the three metrics (FN rate, TC score, SP score) against the nine state-of-the-art tools. We average the scores of the multi-objective solutions in two steps. Since there are 100 solutions in each run, to make the average meaningful, we first sort the 100 solutions according to their scores. Now the scores of the best solutions along all the runs are averaged and taken as the best average score. The same applies for the second best ones and so on. Thus we get the 100 averaged scores for each replicate. Since all replicates actually represent one dataset, we again do a replicate wide averaging in the same way to get the scores of 100 solutions and then plotted them. **Bottom panel** (part (d) - (f)) illustrates how the score of the best solution (within 100 solutions), obtained by the multi-objective formulations, varies across 20 runs. To accomplish this, we collect the best scores (among 100 values) from 20 runs. Thus for each replicate, we get a set of 20 scores. We sort these 20 scores for each replicate and then perform replicate wide averaging (*n*^*th*^ best scores from each replicate are averaged). Finally, we visualize the distribution of the averaged 20 scores using boxplot. In all figures, we depict the averaged scores of nine state-of-the-art tools over 10 replicates using dashed horizontal lines.

The top panel of Figure 2 (part (a) - (c)) illustrates how the 100 solutions provided by the multi-objective formulations perform with respect to each of the three metrics (FN rate, TC score, SP score) against the nine state-ofthe-art tools. We average the scores of the multiobjective solutions in two steps. Since there are 100 solutions in each run, to make the average meaningful, we first sort the 100 solutions according to their scores. Now the scores of the best solutions along all the runs are averaged and taken as the best average score. The same applies for the second best ones and so on. Thus we get the 100 averaged scores for each replicate. Since all replicates actually represent one dataset, we again do a replicate wide averaging in the same way to get the scores of 100 solutions and then plotted them in part (a) to (c) of Figure 2. For the state-of-the-art tools we only needed to average the deterministic score across 10 replicates. We depict these averaged scores using dashed horizontal lines. Moreover, as the 100 solutions are sort along the horizontal axis, it is easy to compare the performance.

In part (a) we notice that {Gap, SOP} and {SimG, SimNG} achieve better FN rates than all state-of-the-art tools. We find PASTA to be the best among the nine tools, followed by T-Coffee. Among the solutions generated by {SimG, SimNG}, on average around, 15% are better than PASTA and 50% are better than T-Coffee. And as for {Gap, SOP}, on average around 10% alignments are better than PASTA and 40% are better than T-Coffee. However, these findings are in contrast with those based on the TC and SP scores (see part (b) and (c)). There we see that the multi-objective formulations barely generate better solutions according to those measures than the best tool (i.e., PASTA).

Next we move to the bottom panel of Figure 2 (part (d) - (f)). Here we illustrate how the score of the best solution (within 100 solutions), obtained by the multi-objective formulations, varies across 20 runs. To do this, we collect the best scores (among 100 values) from 20 runs. Thus for each replicate, we get a set of 20 (best) scores. We sort these 20 scores for each replicate and then performed replicate wise averaging (*n*^*th*^ best scores from each replicate are averaged) similar to what we did earlier. Finally, we visualize the distribution of these averaged 20 scores using boxplots in part (d) to (f) of Figure 2. Here we observe that according to FN rate (part (d)), in case of both objective sets, almost all runs of the metaheuristics produce better solutions than the nine tools. The box width (i.e., interquartile range) suggests that {SimG, SimNG} is slightly more consistent than {Gap, SOP}. But with respect to TC and SP score (part (e) and (f)), we notice that the multi-objective formulations failed to outperform the best tools in significant portion of the cases. Clearly, this can be misleading in the context of phylogeny estimation. Therefore, the experimental results on 100-taxon simulated dataset clearly suggest that, the tools that perform the best based on widely used alignment quality scores are not necessarily the best with respect to phylogenetic tree estimation.

### 2.5 Results on biological rRNA datasets

We conducted ten (instead of 20 as we did on 100-taxon dataset) runs of NSGA-II with our two selected objective sets (i.e., {Gap, SOP} and {SimG, SimNG}) for two datasets, namely, 23S.E and 23S.E.aa_ag.

We compare the performance of the multiobjective formulations with respect to FN rate against the nine state-of-the-art tools from two perspectives in the top panel of Figure 3. Here, part (a) and (b) show the averaged FN rate of 100 solutions over 10 runs. Since each run generates 100 solutions, we make the average meaningful by sorting the 100 FN rates per run. Then we average the best FN rates across all the runs. The same applies for the second best ones and so on. And part (c) and (d) summarize the variation of the best FN rate (among 100 values) across 10 runs. For both of the datasets, FSA performs the best among the nine tools, followed by PRANK for 23.S.E and PASTA for 23S.E.aa ag. The two multi-objective formulations achieved better FN rates than FSA for 23S.E.aa ag (part (b) and (d)). Here, on average, {Gap, SOP} generates around 10% solutions that are better than FSA and 40% solutions that are better than PASTA as shown in part (b). On the other hand, on average {SimG, SimNG} produces very few solutions that are better than FSA but around 40% solutions that are better than PASTA. Part (d) shows that {Gap, SOP} consistently outperforms the best tool (FSA) whereas {SimG, SimNG} outperforms FSA in nearly 40% of the total runs. Now let us see the results for 23S.E (part (a) and (c)), where both of the objective sets remain between the best (FSA) and second best (PRANK) tool. Both of them generates around 5% solutions better than PRANK.

**Figure 3:**
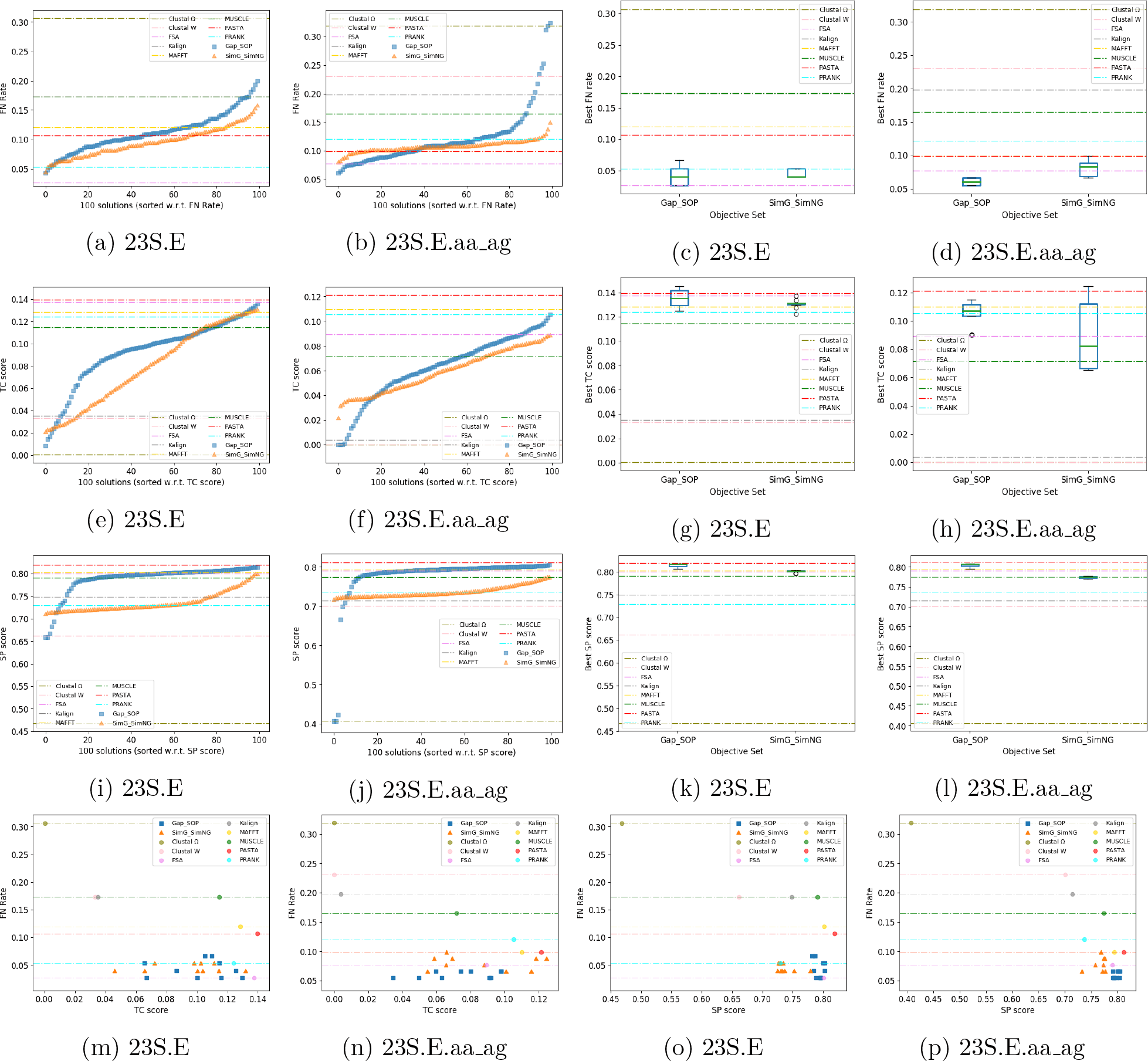
Biological rRNA datasets: **Panel 1 (Top Panel):** part (a) and part (b) show the averaged FN rate of 100 solutions over 10 runs. Since each run generates 100 solutions, we make the average meaningful by sorting the 100 FN rates per run. Then we average the best FN rates along all the runs. The same applies for the second best ones and so on. part (c) and part (d) show the variation of the best FN rates across 10 runs using boxplots. **Panel 2 (Panel 3):** part(e) and part (f) - (part (i) and part (j)) show the TC score (SP score) of 100 solutions averaged over 10 runs. At first, we sort the TC scores (SP scores) of each solution set. Then we average the TC scores (SP score) at each sorted position of all the sets. part (g) and part (h) - (part (k) and part (l)) show the distribution of the best TC scores (SP scores) collected from all runs. **Panel 4 (Bottom panel):** part (m) and part (n) show the relationship between FN rate and TC score for different alignments. part (o) and part (p) show the relationship between FN rate and SP score. In all panels except for the bottom one, we show the performance of nine state-of-the-art tools using dashed horizontal lines; the horizontal lines at the bottom panel mark the FN rates achieved by those tools.

We perform similar analysis based on the widely used two alignment quality measures, namely, TC score and SP score and report the results in Panel 2 and 3 of Figure 3 respectively. We notice that, according to these two popular measures, for both the datasets, the alignments generated by multi-objective formulations failed to beat the best performing tool, PASTA. The clear disagreement between FN rate and TC score (part (m) and (n)) as well as between FN rate and SP score (part (o) and (p)) has been illustrated in Panel 4 of Figure 3. To summarize, from the analysis presented in Panel 4 (bottom panel), we realize that the tools/approaches achieving better performance than our multiobjective formulations in terms of the popular measures, namely, TC score and SP score fail to achieve better FN rates than our multi-objective formulations. To elaborate, according to TC score, PASTA is the best performer among the nine tools, and our objective sets have generated several alignments having worse (lower) TC score than PASTA (and FSA). However those alignments can produce phylogenetic trees with better FN rates than those tools. Even from among the tools, there is disagreement between TC score and FN rate: PASTA is in fact behind FSA in terms of the latter. Similarly in part (o) and (p) of Panel 4 which is dedicated to the comparison between FN rate and SP score, we find several alignments generated by the multi-objective formulations that are worse than PASTA in terms of SP score, but, achieve better FN rates than that tool.

### 2.6 Results on BAliBASE datasets

For each of the selected BAliBASE datasets under six groups (RV11, RV12, RV20, RV30, RV40 and RV50), we conducted 20 independent runs of NSGA-II considering its stochastic nature according to the standard practice of OR literature. Once again we analyze the generated solutions based on the quality of alignments and resultant trees. We witnessed that the alignments that are better according to widely accepted alignment scores, not necessarily generate better phylogenetic trees. Here we discuss our key observations on the selected four datasets (BB11005, BB11018, BB11020 and BB11033) under the group RV11. For the remaining groups (RV12, RV20, RV30, RV40 and RV50), our obtained results are similar. For the sake of brevity, we present those results in Section S3 of the supplementary file. Figures 5 and 4 present the results of our experiments on the datasets of group RV11. According to FN rate (part (a) - (h) of Figure 5), at least one of the two objective sets generates better or equivalent solutions than the best tool throughout all the instances. For BB11020, {SimG, SimNG} can achieve 12% FN rate as opposed to 50% FN rate attained by the best tool which is a huge improvement. Considering TC score (part (a) - (h) of Figure 4), the two objective sets can outperform all the tools only for BB11020 which is contrary to the findings based on FN rate. So again we see the disagreement between FN rate and TC score which we examine graphically in part (i) - (l) of Figure 5. If we observe the results based on SP score (part (i) - (p) of Figure 4), we get similar disagreement between FN rate and SP score which is illustrated in part (m) - (p) of Figure 5. These figures provide evidence that a solution with the best TC and/or SP score may not give the best FN rate.

**Figure 4:**
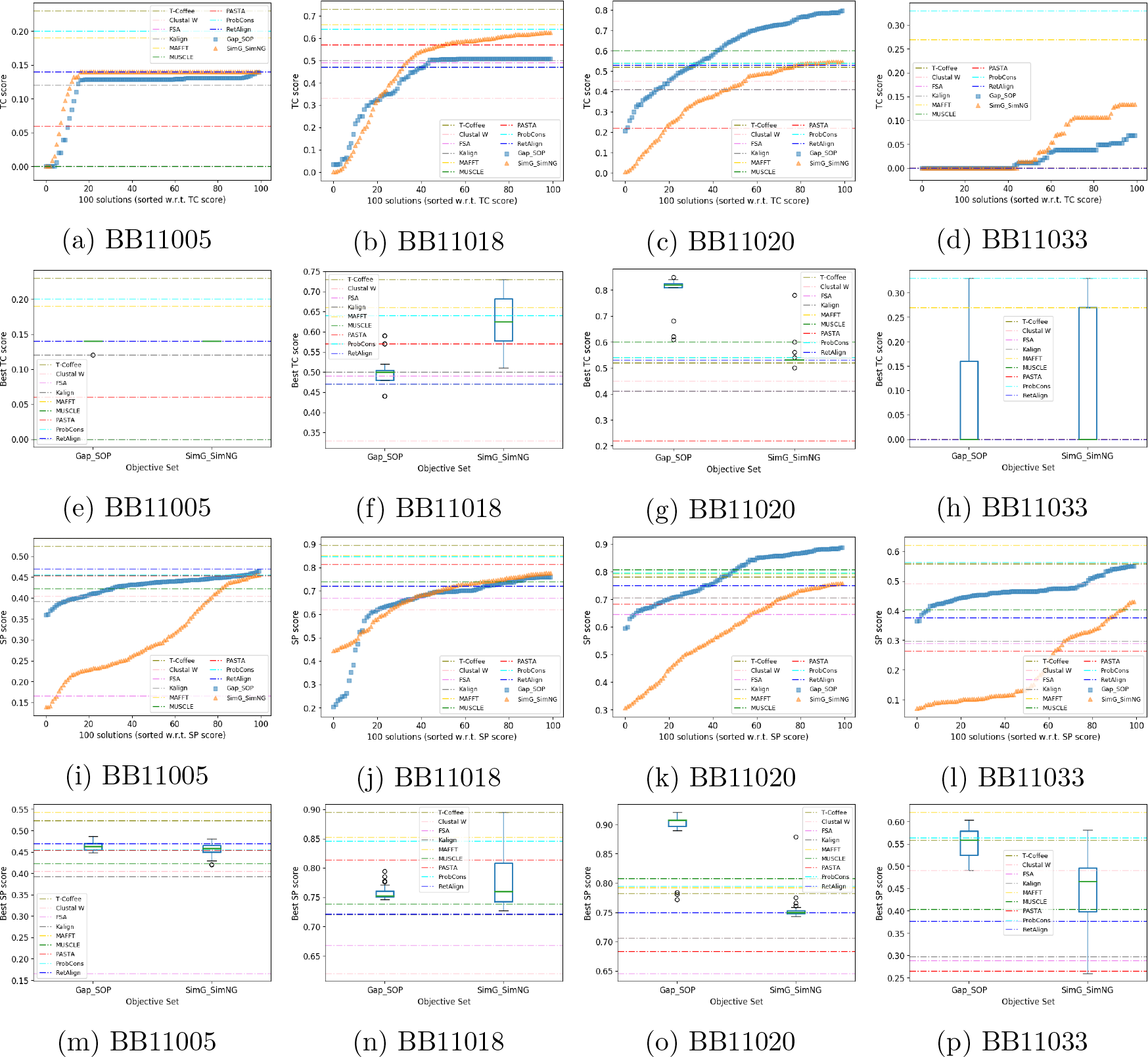
RV11: **Panel 1 (Panel 3)**: part (a) - (d) - (part (i) - (l)) shows the TC score (SP score) of 100 solutions averaged over 20 runs. At first, we sort the TC scores (SP scores) of each solution set. Then we average the TC scores (Sp scores) at each sorted position of all the sets. **Panel 2 (Panel 4)**: part (e) - (h) - (part (m) - (p)) shows the distribution of the best TC scores (Sp scores) collected from all runs. In all panels, we show the performance of nine state-of-the-art tools using dashed horizontal lines.

**Figure 5:**
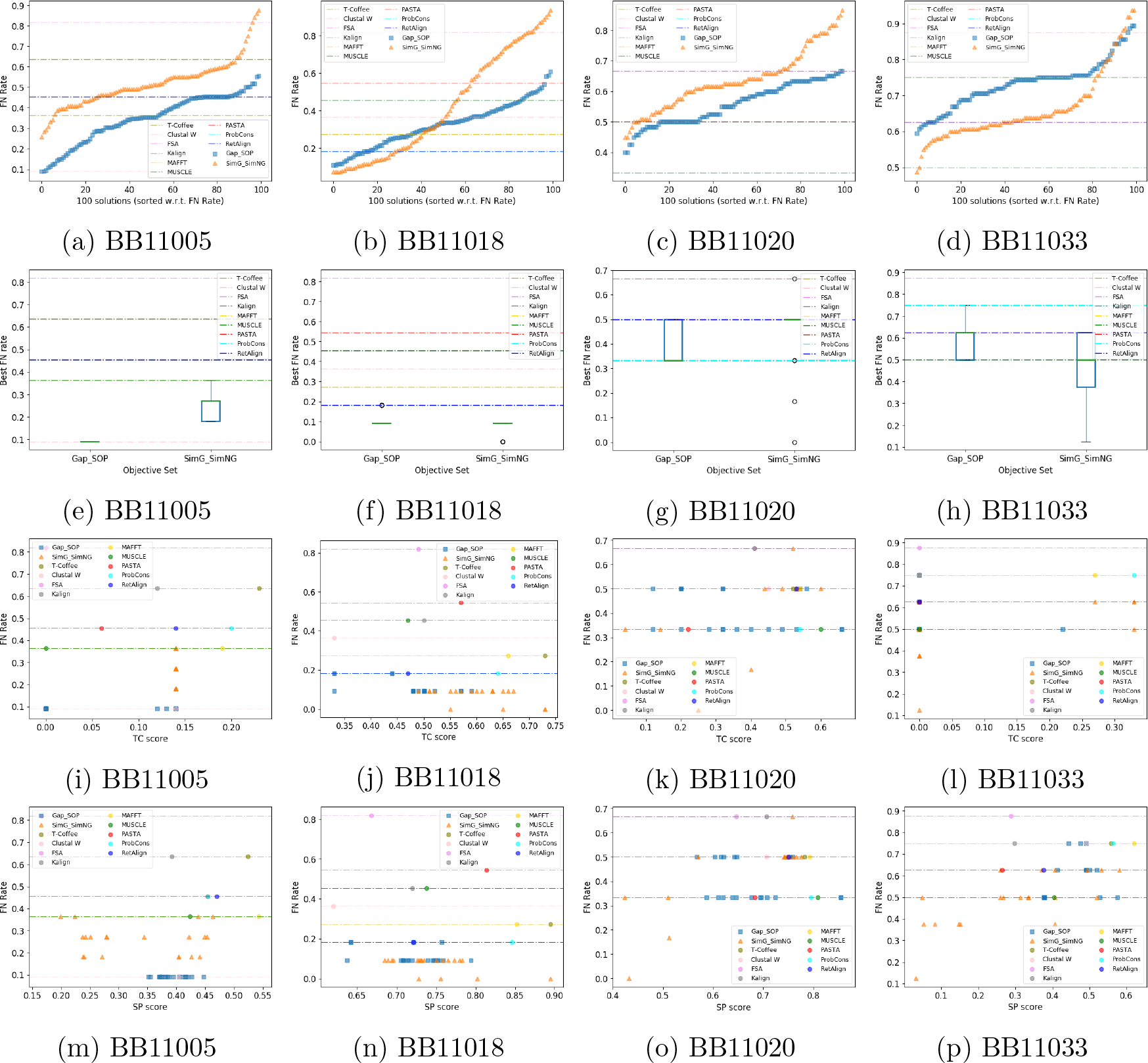
RV11. **Panel 1 (Top panel):**part (a) - (d) show the FN rate of 100 solutions averaged over 20 runs. At first, we sort the FN rates of each solution set. Then we average the FN rates at each sorted position of all the sets. **Panel 2:**part (e) - (h) show the distribution of the best FN rates collected from all runs. **Panel 3 (Panel 4):**part (i) - (l) - (part (m) - (p)) show the relationship between FN rate and TC score (SP score) for different alignments. In all panels, we show the FN rates achieved by the nine state-of-the-art tools using dashed horizontal lines.

#### 2.6.1 Statistical significance

Now we confirm the significance of the improvement achieved by the multi-objective formulations in terms of FN rate over nine MSA tools on 27 BAliBASE datasets by applying an appropriate statistical test. We form paired data by picking the FN rate achieved by each (MSA method, dataset) pair. For the metaheuristics, we take the average of the 20 best FN rates from 20 independent runs considering its stochastic nature. As our data do not satisfy the condition of normality and homoscedasticity [36], we choose a series of nonparametric tests following the recommendation of [37]. At first, we simultaneously compare all the methods using the Friedman test [38] which gives the relative ranking (lower is better) of all the methods and strongly suggests the existence of significant differences among the methods considered (as *p*-value is 0). The results have been presented in Column 2 of Table 1. Here we see that the multi-objective formulations achieve the top two positions. Next, we complement the Friedman test by following Holm’s post-hoc procedure [39] to contrast the difference between the multi-objective formulations and each of the nine tools. The results have been summarized in Columns 3 and 4 of Table 1. Here, each cell shows the adjusted *p*-value which indicates the significance of difference in performance (based on FN rate) between two methods. We notice that all the *p*-values are very close to 0 and the values for {SimG, SimNG} are lower than {Gap, SOP}. So we can state with high confidence that, the multi-objective formulations achieve statistically significant improvement over the nine MSA tools.

**Table 1:** Friedman test (Column 2) The Average Friedman’s ranking (lower is better) achieved by the MSA methods over 27 BAliBASE datasets. We performed the Friedman test based on FN rate achieved by the tools. For our metaheuristics based multi-objective formulations, we consider the average of the 20 best FN rates obtained from 20 runs. We also show the computed statistics and corresponding *p*-value. <u>Holm’s post-hoc procedure (Columns 3 and 4):</u> Comparison between the metaheuristics and the MSA tool using the Holm’s post-hoc procedures (as a complement of the Friedman test) over 27 BAliBASE datasets. Each entry shows the adjusted *p*-value which indicates the significance of difference in performance (based on FN rate) between two methods.

| 1 | 2 | 3 | 4 |
| --- | --- | --- | --- |
| Method | Friedman’s Rank* | Holm’s adjusted $p$-value | |
|  |  | NSGA-II<sub>{SimG, SimNG}</sub> | NSGA-II<sub>{Gap, SOP}</sub> |
| NSGA-II<sub>{SimG, SimNG}</sub> | 2.2037 | - | 0.46018 |
| NSGA-II<sub>{Gap, SOP}</sub> | 2.8704 | 0.46018 | - |
| ProbCons | 5.7963 | 0.00014 | 0.00238 |
| Clustal $\Omega$ | 6.2963 | 0.00002 | 0.00044 |
| MAFFT | 6.4074 | 0.00001 | 0.00036 |
| Kalign | 6.7037 | 0.00000 | 0.00011 |
| PASTA | 6.8148 | 0.00000 | 0.00007 |
| FSA | 6.9444 | 0.00000 | 0.00004 |
| MUSCLE | 7.1482 | 0.00000 | 0.00002 |
| Clustal W | 7.3519 | 0.00000 | 0.00001 |
| RetAlign | 7.4630 | 0.00000 | 0.00000 |
| *Statistic | 10.5911 | N/A |  |
| *$p$-value | 0.00000 | | |

## 3 Discussion

In this study, we have introduced a phylogenyaware multi-objective optimization approach to compute MSA with an ultimate goal to infer the phylogenetic tree from the resultant alignments. To optimize MSA, we proposed two simple objective functions in addition to the existing ones. We judged the potential capability of each objective function to yield better trees by employing domain knowledge as well as by applying statistical approaches. We employed multiple linear regression to measure the degree of association between the individual objective functions and the quality of inferred phylogenetic tree (i.e., FN rate). Thus, we provide empirical justification to choose two multi-objective formulations to move forward. Afterwards, we performed extensive experimentation with both simulated and biological datasets to demonstrate the benefit of our approach. We showed that the simultaneous optimization of a set of phylogeny-aware objective functions can lead to phylogenetic trees with improved accuracy than that of the state-of-theart MSA tools. From this finding, we would like to hypothesize that, the use of domain specific measures can aid an MSA methods in other application domain as well.

Standard criteria (SP-score, TC-score, etc.) for assessing alignment quality are usually based on shared homology pairs (SP score) or identical columns (TC score), and do not explicitly consider a particular application domain. Mistakes in alignments that are not important with respect to an application domain may not impact the ultimate accuracy of that particular inference. For example, not all sites are significant with respect to protein structure and function prediction, and hence multiple alignments with different accuracy may lead to the same predictions [1]. Similarly, in the context of phylogeny estimation, alignments with substantially different SP scores may lead to trees with the same accuracy [3]. In this study, we systematically investigate the impact of evaluation criteria of an alignment on phylogenetic tree inference problem. Our results suggest that it could be possible to develop improved MSA methods for phylogenetic analysis by carefully choosing appropriate objective functions. Moreover, in almost all existing studies on MSA, we find the researchers evaluating the effectiveness of MSA methods using some generic alignment quality measures (i.e., TC score, SP score). Contrastingly, our results revealed that optimizing those widely used measures do not necessarily lead us to the best phylogenetic tree. This finding could be an eye opener for the researchers who need to use MSA methods to address a particular application.

Our findings and proposed multi-objective formulation can be particularly beneficial for iterative methods like SATé and PASTA that iteratively co-estimate both alignment and tree. These methods obtain an initial alignment and a tree that guide each other to improved estimates in an iterative fashion. They make an effort to exploit the close association between the accuracy of an MSA and the corresponding tree in finding the output through multiple iterations from both directions. Therefore, carefully choosing an evaluation metric for an MSA with better correlation to the tree accuracy seems likely to improve the results of these co-estimation techniques. Thus, our methodology, if adopted, may potentially have a profound positive impact on the accuracy of these iterative co-estimation techniques.

This study will encourage the scientific community to investigate various application-aware measures for computing and evaluating MSAs. This will potentially prompt more experimental studies addressing specific application domains; and ultimately will propel our understanding of MSAs and their impact in various domains in computational biology, i.e, phylogeny estimation, protein structure and function prediction, orthology prediction etc. This study will also encourage the researchers to develop new scalable MSA tools by simultaneously optimizing multiple appropriate optimization criteria. Thus, we believe that this study will pioneer new models and optimization criteria for computing MSA – laying a firm, broad foundation for application specific multi-objective formulation for estimating multiple sequence alignment.

We performed an extensive experimental study comprising 29 datasets of varying sizes and complexities, and our findings are consistent throughout all the datasets. Still, we acknowledge the possibility of facing few unforeseen circumstances as follows. There might be some datasets on which our approach might not exhibit satisfactory performance. Besides, currently we did not pay any effort to improve the running time of our approach which is higher as compared to top MSA tools. However, sufficient speedup could be achieved by leveraging the modern computing architectures (computer cluster, GPU, etc.).

Formulating phylogeny-aware multi-objective formulation (application specific evaluation criteria in general) cannot be developed entirely in one study; it should evolve in response to scientific findings and systematists’ feedback. This requires active involvement of evolutionary biologists, computer scientists, systematists, and others – leading to improved understandings of alignments and how they are related to various fields in comparative genomics.

## 4 Methods

We begin this section by introducing the objective functions that we used to perform MSA. Then, we present the multi-objective metaheuristics selected to optimize those objective functions. Afterwards, we describe our method of evaluating estimated alignments and phylogenetic trees. Finally, we summarize the methodology followed to establish the effectiveness of our objectives.

### 4.1 Objective functions

Most real-world optimization problems naturally work towards achieving several objectives. Some of these objectives are conflicting to each other. However, these problems can be transformed into single-objective ones using various simplifying techniques to avoid complexities [34]. On the contrary, a multi-objective formulation defines the problem using a set of objective functions and subsequently specialized methods can be applied to optimize all the objectives simultaneously. In this study, we have selected the following three multi-objective formulations of MSA from the literature based on their simplicity as well as performance as reported in the literature.

- {SOP, TC}: Maximize the sum of pairs (SOP) and the number of totally aligned columns (TC) [20].
- {Gap, SOP}: Maximize the sum of pairs (SOP) and minimize the number of gaps (Gap) [23].
- {wSOP, TC}: Maximize the weighted sum of pairs with affine gap penalties (wSOP) and the number totally aligned columns (TC) [31, 26].

We describe these objective functions along with several existing ones in Section S1 of the supplementary file. Now, we propose four new objective functions that quantify different aspects of an MSA. Unlike the existing objective functions in the literature, we avoid combining multiple aspects of an MSA into a single objective. We introduce them as follows:

- **Minimize entropy (Entropy)**: We modify the usual definition by considering only non-gap column for the calculation of entropy.
- **Maximize similarity based on gap containing columns (SimG)**: Here we calculate similarity only for those columns that contain at least one gap.
- **Maximize similarity based on non-gap columns (SimNG)**: We consider only nongap column while measuring similarity.
- **Maximize concentration of gaps (GapCon)**: We find that a widely used objective function, namely, affine gap penalty [25], combines two aspects of an aligned sequence, number of gaps and concentration of gaps, into a single one using weighted sum. We need to tune the weight values based on the dataset. To avoid this tuning we decide to decouple the two components. We have already considered the number of gaps as an objective function. Now we define the concentration of gaps as an independent objective which as calculated as follows. For each sequence, we count the number of consecutive gaps and take the mean of these counts. Finally, we average the resultant means for all sequences.

In this study, we used the terms shown in Table 2 to refer to these objective functions.

We need a substitution matrix to calculate SOP and wSOP [40]. The values of this substitution matrix depend on the trait of a particular dataset. In this study, we used NUC4.4 (supplied by NCBI at ftp://ftp.ncbi.nih.gov/blast/matrices/NUC.4.4) for nucleotide sequences and BLOSUM62 [41] for protein sequences. On the contrary, the four objective functions that we proposed are non-parametric which are independent of the dataset.

### 4.2 Multi-objective metaheuristics

To simultaneously optimize multiple objective functions, we ran two popular multi-objective metaheuristics: NSGA-II [35] and NSGAIII [32]. They belong to the class of evolutionary algorithms. Several studies ([27, 21, 42]) demonstrated the strength of NSGA-II for solving MSA. NSGA-II works best when the number of objectives is upto three while NSGA-III is specially designed for handling more than three objectives. Hence, we applied these algorithms according to Table 3. We discuss these methods along with their vital components in Section S2 of the supplementary file.

**Table 3:**
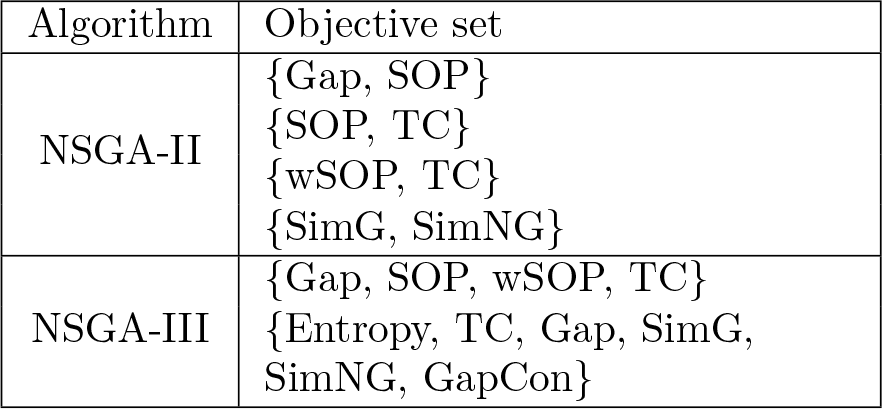
Our selected algorithms and corresponding objective set.

| Algorithm | Objective set |
| --- | --- |
| NSGA-II | {Gap, SOP}<br>{SOP, TC}<br>{wSOP, TC}<br>{SimG, SimNG} |
| NSGA-III | {Gap, SOP, wSOP, TC}<br>{Entropy, TC, Gap, SimG, SimNG, GapCon} |

### 4.3 State-of-the-art MSA tools

We used the alignments generated by nine representative state-of-the-art MSA tools (shown in Table 4) to compare with our approach. We run each of them with its default parameter configuration. Moreover, we initialize the multiobjective metaheuristics with a set of alignments generated by randomly mixing and modifying those nine alignments. Notably, this approach, known as the seeded initial population generation, is quite common in the metaheuristics literature specially for multi-objective optimization.

**Table 4:** List of state-of-the-art MSA tools that we used in this study.

| For nucleotide sequences |  | For protein sequences |  |
| --- | --- | --- | --- |
| Tool | Version | Tool | Version |
| FSA [7] | 1.15.9 | FSA | 1.15.9 |
| PASTA [2] | 1.7.8 | PASTA | 1.7.8 |
| T-Coffee [9] | 11.00 | T-Coffee | 11.00 |
| MAFFT [13] | 7.31 | MAFFT | 7.245 |
| Clustal W [43] | 2.1 | Clustal W | 2.1 |
| Clustal $\Omega$ [4] | 1.2.4 | RetAlign [8] | 1.0 |
| MUSCLE [14] | 3.8.31 | MUSCLE | 3.8.31 |
| PRANK [5] | 0.170427 | ProbCons [10] | 1.12 |
| Kalign [6] | 2.03 | Kalign | 2.04 |

### 4.4 Evaluation of estimated alignments

We evaluate estimated alignments with respect to reference alignment using two well-known alignment quality scores called TC score and SP score. These two scores are defined below:

- TC score is the ratio of the number of correctly aligned columns in the estimated alignment to the total number of aligned columns in the reference alignment. This is also known as column score.
- SP score is the ratio of the number of aligned pairs in the estimated alignment to the total number of aligned pairs in the reference alignment.

For both the measures, higher value implies better score.

### 4.5 Phylogenetic tree estimation

For each of the generated alignment we estimate the phylogenetic tree using Maximum Likelihood (ML) method. We used a popular tool named FastTree-2 developed by 44.

### 4.6 Evaluation of phylogenetic tree

We evaluate the quality of each estimated ML tree with respect to the true phylogenetic tree using a widely used measure known as the False Negative (FN) rate. FN rate is the percentage of edges present in the true tree but missing in the estimated tree. So a small value of FN rate is desirable.

### 4.7 Evaluation of objective functions

In the context of phylogeny estimation, a desired objective function for MSA should lead to such alignments which can produce highly accurate (having small FN rate) ML trees. Considering this fact, we try to evaluate the effectiveness of an objective function by studying how its values are associated with the corresponding FN rates. The objective function that frequently exhibits positive correlation with FN rate is predicted to be a good optimization criteria. To accomplish this, we fit multiple linear regression model to calculate the degree of association (i.e., regression coefficient) between an objective and FN rate. Then we apply t-test, with null hypothesis that there is no association, to check the significance of individual regression coefficients. It should be noted that, such regression results does not necessarily indicate the strength of an objective as an optimization criterion. However, such results can definitely be utilized as the starting point for experimentation for further validation.

## Author Contributions

**MAN** conceived the study, implemented the methods, designed and conducted the experiments, analyzed and interpreted the results and wrote the first draft; **MSB** conceived the study, designed and supervised the experiments, selected the datasets, analyzed and interpreted the results; **AHR** designed the statistical tests and analyzed and interpreted the results; **RS** checked the implementation, analyzed and interpreted the results and co-supervised the overall research work; **MSR** conceived the study, designed and supervised the experiments, analyzed and interpreted the results and supervised the overall research work. **All authors** took part in finalizing the draft and approved the final manuscript.

## Acknowledgements

**MAN** is a recipient of the ICT PhD Fellowship administered by ICT Division, Government of People’s Republic of Bangladesh.

## Competing interests

The authors declare no competing interests.

## References

[1] Tandy Warnow. Large-scale multiple sequence alignment and phylogeny estimation. In Models and algorithms for genome evolution, pages 85–146. Springer, 2013.

[2] Siavash Mirarab, Nam Nguyen, Sheng Guo, Li-San Wang, Junhyong Kim, and Tandy Warnow. Pasta: ultra-large multiple sequence alignment for nucleotide and aminoacid sequences. Journal of Computational Biology, 22(5):377–386, 2015.

[3] Kevin Liu, Sindhu Raghavan, Serita Nelesen, C Randal Linder, and Tandy Warnow. Rapid and accurate large-scale coestimation of sequence alignments and phylogenetic trees. Science, 324(5934):1561–1564, 2009.

[4] Fabian Sievers, Andreas Wilm, David Dineen, Toby J Gibson, Kevin Karplus, Weizhong Li, Rodrigo Lopez, Hamish McWilliam, Michael Remmert, Johannes Söding, et al. Fast, scalable generation of high-quality protein multiple sequence alignments using clustal omega. Molecular systems biology, 7(1):539, 2011.

[5] Ari Löoytynoja and Nick Goldman. An algorithm for progressive multiple alignment of sequences with insertions. Proceedings of the National academy of sciences of the United States of America, 102(30):10557–10562, 2005.

[6] Timo Lassmann, Oliver Frings, and Erik LL Sonnhammer. Kalign2: high-performance multiple alignment of protein and nucleotide sequences allowing external features. Nucleic acids research, 37(3):858–865, 2008.

[7] Robert K Bradley, Adam Roberts, Michael Smoot, Sudeep Juvekar, Jaeyoung Do, Colin Dewey, Ian Holmes, and Lior Pachter. Fast statistical alignment. PLoS computational biology, 5(5):e1000392, 2009.

[8] Adrienn Szabö, Ádáam Novák, István Miklós, and Jotun Hein. Reticular alignment: A progressive corner-cutting method for multiple sequence alignment. BMC bioinformatics, 11(1):570, 2010.

[9] Cédric Notredame, Desmond G Higgins, and Jaap Heringa. T-coffee: a novel method for fast and accurate multiple sequence alignment1. Journal of molecular biology, 302(1):205–217, 2000.

[10] Chuong B Do, Mahathi SP Mahabhashyam, Michael Brudno, and Serafim Batzoglou. Probcons: Probabilistic consistency-based multiple sequence alignment. Genome research, 15(2):330–340, 2005.

[11] Yongchao Liu, Bertil Schmidt, and Douglas L Maskell. Msaprobs: multiple sequence alignment based on pair hidden markov models and partition function posterior probabilities. Bioinformatics, 26(16):1958–1964, 2010.

[12] Usman Roshan and Dennis R Livesay. Probalign: multiple sequence alignment using partition function posterior probabilities. Bioinformatics, 22(22):2715–2721, 2006.

[13] Kazutaka Katoh, Kazuharu Misawa, Keiichi Kuma, and Takashi Miyata. Mafft: a novel method for rapid multiple sequence alignment based on fast fourier transform. Nucleic acids research, 30(14):3059–3066, 2002.

[14] Robert C Edgar. Muscle: multiple sequence alignment with high accuracy and high throughput. Nucleic acids research, 32(5):1792–1797, 2004.

[15] Jimin Pei and Nick V Grishin. Mummals: multiple sequence alignment improved by using hidden markov models with local structural information. Nucleic acids research, 34(16):4364–4374, 2006.

[16] Shinsuke Yamada, Osamu Gotoh, and Hayato Yamana. Improvement in accuracy of multiple sequence alignment using novel group-to-group sequence alignment algorithm with piecewise linear gap cost. BMC bioinformatics, 7(1):524, 2006.

[17] Cédric Notredame and Desmond G Higgins. Saga: sequence alignment by genetic algorithm. Nucleic acids research, 24(8):1515–1524, 1996.

[18] Álvaro Rubio-Largo, Leonardo Vanneschi, Mauro Castelli, and Miguel A VegaRodríguez. A characteristic-based framework for multiple sequence aligners. IEEE transactions on cybernetics, 48(1):41–51, 2018.

[19] Julie D Thompson, Benjamin Linard, Odile Lecompte, and Olivier Poch. A comprehensive benchmark study of multiple sequence alignment methods: current challenges and future perspectives. PloS one, 6(3):e18093, 2011.

[20] Fernando José Mateus da Silva, Juan Manuel Sánchez Pérez, Juan Antonio Gómez Pulido, and Miguel A Vega Rodríguez. Alineagaa genetic algorithm with local search optimization for multiple sequence alignment. Applied Intelligence, 32(2):164–172, 2010.

[21] Francisco M Ortunño, Olga Valenzuela, Fernando Rojas, Hector Pomares, Javier P Florido, Jose M Urquiza, and Ignacio Rojas. Optimizing multiple sequence alignments using a genetic algorithm based on three objectives: structural information, non-gaps percentage and totally conserved columns. Bioinformatics, 29(17):2112–2121, 2013.

[22] Wilson Soto and David Becerra. A multiobjective evolutionary algorithm for improving multiple sequence alignments. In Brazilian Symposium on Bioinformatics, pages 73–82. Springer, 2014.

[23] Maryam Abbasi, Luís Paquete, and Francisco B Pereira. Local search for multiobjective multiple sequence alignment. In International Conference on Bioinformatics and Biomedical Engineering, pages 175–182. Springer, 2015.

[24] Huazheng Zhu, Zhongshi He, and Yuanyuan Jia. A novel approach to multiple sequence alignment using multiobjective evolutionary algorithm based on decomposition. IEEE journal of biomedical and health informatics, 20(2):717–727, 2016.

[25] R Ranjani Rani and D Ramyachitra. Multiple sequence alignment using multiobjective based bacterial foraging optimization algorithm. Biosystems, 150:177–189, 2016.

[26] Álvaro Rubio-Largo, Miguel A VegaRodríguez, and David L González-Álvarez. A hybrid multiobjective memetic metaheuristic for multiple sequence alignment. IEEE Transactions on Evolutionary Computation, 20(4):499–514, 2016.

[27] Cristian Zambrano-Vega, Antonio J Nebro, José García-Nieto, and José F Aldana-Montes. Comparing multi-objective metaheuristics for solving a three-objective formulation of multiple sequence alignment. Progress in Artificial Intelligence, pages 1–16, 2017.

[28] Alexandros Stamatakis. Raxml version 8: a tool for phylogenetic analysis and postanalysis of large phylogenies. Bioinformatics, 30(9):1312–1313, 2014.

[29] Siavash Mirarab and Tandy Warnow. Fastsp: linear time calculation of alignment accuracy. Bioinformatics, 27(23):3250–3258, 2011.

[30] Julie D Thompson, Patrice Koehl, Raymond Ripp, and Olivier Poch. Balibase 3.0: latest developments of the multiple sequence alignment benchmark. Proteins: Structure, Function, and Bioinformatics, 61(1):127–136, 2005.

[31] Álvaro Rubio-Largo, Miguel A Vega-Rodríguez, and David L GonzálezÁAlvarez. Hybrid multiobjective artificial bee colony for multiple sequence alignment. Applied Soft Computing, 41:157–168, 2016.

[32] Kalyanmoy Deb and Himanshu Jain. An evolutionary many-objective optimization algorithm using reference-point-based nondominated sorting approach, part i: Solving problems with box constraints. IEEE Trans. Evolutionary Computation, 18(4):577–601, 2014.

[33] Douglas C Montgomery, Elizabeth A Peck, and G Geoffrey Vining. Introduction to linear regression analysis, volume 821. John Wiley & Sons, 2012.

[34] Deb Kalyanmoy. Multi objective optimization using evolutionary algorithms. John Wiley and Sons, 2001.

[35] Kalyanmoy Deb, Amrit Pratap, Sameer Agarwal, and TAMT Meyarivan. A fast and elitist multiobjective genetic algorithm: Nsga-ii. IEEE transactions on evolutionary computation, 6(2):182–197, 2002.

[36] David J Sheskin. Handbook of parametric and nonparametric statistical procedures. crc Press, 2003.

[37] Joaquín Derrac, Salvador García, Daniel Molina, and Francisco Herrera. A practical tutorial on the use of nonparametric statistical tests as a methodology for comparing evolutionary and swarm intelligence algorithms. Swarm and Evolutionary Computation, 1(1):3–18, 2011.

[38] Milton Friedman. The use of ranks to avoid the assumption of normality implicit in the analysis of variance. Journal of the american statistical association, 32(200):675–701, 1937.

[39] Sture Holm. A simple sequentially rejective multiple test procedure. Scandinavian journal of statistics, pages 65–70, 1979.

[40] Richard Durbin, Sean R Eddy, Anders Krogh, and Graeme Mitchison. Biological sequence analysis: probabilistic models of proteins and nucleic acids. Cambridge university press, 1998.

[41] Steven Henikoff and Jorja G Henikoff. Amino acid substitution matrices from protein blocks. Proceedings of the National Academy of Sciences, 89(22):10915–10919, 1992.

[42] Cristian Zambrano-Vega, Antonio J Nebro, José García-Nieto, and Jose F Aldana-Montes. M2align: parallel multiple sequence alignment with a multiobjective metaheuristic. Bioinformatics, 33(19):3011–3017, 2017.

[43] Julie D Thompson, Desmond G Higgins, and Toby J Gibson. Clustal w: improving the sensitivity of progressive multiple sequence alignment through sequence weighting, position-specific gap penalties and weight matrix choice. Nucleic acids research, 22(22):4673–4680, 1994.

[44] Morgan N Price, Paramvir S Dehal, and Adam P Arkin. Fasttree 2-approximately maximum-likelihood trees for large alignments. PloS one, 5(3):e9490, 2010.

